# A candidate neuroimaging biomarker for detection of neurotransmission-related functional alterations and prediction of pharmacological analgesic response in chronic pain

**DOI:** 10.1101/2021.02.17.431572

**Authors:** Daniel Martins, Mattia Veronese, Federico Turkheimer, Matthew A. Howard, Steven CR Williams, Ottavia Dipasquale

## Abstract

**Background:** Chronic pain is a world-wide clinical challenge. Response to analgesic treatment is limited and difficult to predict. Functional MRI (fMRI) has been suggested as a potential solution. However, while most analgesics target specific neurotransmission pathways, fMRI-based biomarkers are not specific for any neurotransmitter system, limiting our understanding of how they might contribute to predict treatment response.

**Methods:** Here, we sought to bridge this gap by applying Receptor-Enriched Analysis of Functional Connectivity by Targets (REACT) to investigate whether neurotransmission-enriched functional connectivity (FC) mapping can provide insights into the brain mechanisms underlying chronic pain and inter-individual differences in analgesic response after a placebo or duloxetine. Chronic knee osteoarthritis (OA) pain patients (n=56) underwent pre-treatment brain scans in two clinical trials. Study 1 (*n*=17) was a 2-week single-blinded placebo pill trial. Study 2 (*n*=39) was a 3-month double-blinded randomized trial comparing placebo to duloxetine, a dual serotonin-noradrenaline reuptake inhibitor.

**Results:** Across two independent studies, we found that chronic pain OA patients present FC alterations in the FC related to the serotonin (SERT) and noradrenaline (NET) transporters, when compared to age-matched healthy controls. Placebo responders presented with higher pre-treatment dopamine transporter (DAT)-enriched FC than non-responders. Duloxetine responders presented with higher pre-treatment SERT and NET-enriched FC than non-responders. Pre-treatment SERT and NET-enriched FC achieved predictive positive values of duloxetine response up to 85.71%.

**Conclusion:** Neurotransmission-enriched FC mapping might hold promise as a new mechanistic-informed biomarker for functional brain alterations and prediction of response to pharmacological analgesia in chronic pain.

## Introduction

Pain is a world-wide leading cause of disability, constituting one of the primary reasons for people to seek healthcare(1–3). Chronic pain is a disease in its own right, characterized by persistence of pain beyond normal healing time(1). Despite the high personal and societal costs(4), pain management in patients with chronic pain is still generally unsatisfactory(5). Although the number of potential treatments has grown substantially (i.e. antidepressants, anticonvulsants, opioids)(6), treatment response is overall low(7) and why only some patients respond remains poorly understood(8). On the other, most of the available pharmacological treatments for patients with chronic pain are accompanied by considerable side effects and risk of misuse (i.e. opioids)(9), motivating high rates of treatment non-adherence(10). A strong case has been made for a mechanism-based and individualized approach to chronic pain therapy(11); yet, our capacity to predict who may or may not benefit from a specific analgesic treatment is still limited(12), leading high numbers of non-responsive patients to experience a range of side effects with minimal or null clinical benefit. Therefore, developing mechanism-based biomarkers that can guide analgesic treatment selection for chronic pain patients based on prediction of treatment response remains an unmet target and a clinical need.

Part of this problem stems from our limited understanding of the neurobiological mechanisms underlying chronic pain and, hence, of the mechanisms through which most of these pharmacological treatments might produce persistent pain relief in chronic pain patients(12). Currently, it is generally accepted that chronic pain is a multifactorial entity entailing physical, psychological, emotional, and social aspects(1). Preclinical studies have offered insights into key central mechanisms that might contribute to chronic pain, including sensitization phenomena in an array of nervous system pathways, imbalances in the facilitatory and inhibitory descending modulation pathways from the brain that regulate the transmission of noxious information in the spinal cord, neuroinflammation and glial dysfunction, among others(13–17). These findings have fuelled substantial interest in developing neuroimaging-based biomarkers that could unravel how chronic pain affects brain functioning and what form of brain pathophysiology in these patients can be targeted by different treatments(18, 19). While a range of preliminary diagnosis, prognosis and treatment response brain biomarkers have been suggested (for extensive reviews please see (18, 19)), to date these biomarkers have provided minimal direct clinical application in the management of chronic pain patients.

Most pharmacological analgesic treatments target specific neurotransmission pathways. For instance, duloxetine, a dual antidepressant often used to manage pain in chronic pain patients, inhibits the reuptake of both serotonin and noradrenaline, increasing their bioavailability in the synapses(20). However, neuroimaging-based biomarkers of brain function (i.e. such as those based on measurements of BOLD signal(21)) are not specific for any neurotransmission system, limiting the potential mechanistic understanding of how these biomarkers might contribute to explain treatment response. The same limitation applies to the potential neuroimaging biomarkers in unravelling functional changes in neuromodulatory pathways that could guide drug development or repurposing for patients with chronic pain(12).

Here, we sought to bridge this gap by applying the recently developed Receptor-Enriched Analysis of Functional Connectivity by Targets (REACT) multimodal framework(22), which enriches resting-state functional magnetic resonance imaging (rs-fMRI) analysis with information about the distribution density of molecular targets derived from PET and SPECT imaging(22, 23). We performed secondary analyses of two openly available rs-fMRI datasets(24) to investigate two main questions: i) do patients with painful chronic knee osteoarthritis present functional alterations in key neurotransmitter-related circuits associated with pain control and regulation, when compared to age-matched healthy controls? ii) Can pre-treatment inter-individual differences in the functional connectivity (FC) of these neurotransmitter-related circuits predict analgesia in response to placebo or treatment with duloxetine? Chronic knee osteoarthritis (OA) pain patients (*n* = 56) underwent pre-treatment brain scans in two clinical trials. Study 1 (*n* = 17) was a 2-week single-blinded placebo pill trial. Study 2 (*n* = 39) was a 3-month double-blinded, between-subject, randomized trial comparing placebo (n = 20) to duloxetine (n = 19). Patients were compared to healthy controls (n=20).

We focused our analyses on the functional circuits related to the serotonin (SERT), noradrenaline (NET) and dopamine (DAT) transporters, and the μ-opioid receptor, as general indicators of the regional distribution of the neurotransmission related to the serotonin, noradrenaline, dopamine and opioid systems, respectively. We informed our selection of molecular targets by the fact that these neurotransmitter systems play pivotal roles in pain regulation, namely in those descendent modulatory pathways controlling the spinal transmission of nociceptive information(25, 26). Furthermore, they have also been implicated in placebo analgesia (i.e. dopamine and opioids)(27) and correspond to the main molecular targets of duloxetine (i.e. serotonin and noradrenaline transporters)(20), which treatment response was effectively studied herein.

## Results

Here, we report in detail only the results of our analyses on molecular-enriched FC and summarize below some of the main findings from the original analyses(24) on sociodemographic and clinical variables that might help to interpret our novel imaging findings. For a detailed description of sociodemographic and clinical variables we refer the reader to the original article published elsewhere(24).

In study 1, eight patients met the criteria for placebo response and nine patients were classified as non-responders. Responders and non-responders did not differ in baseline pain ratings, age, disease duration, depressive symptoms, pain catastrophizing or medication use at the entry of the study. In study 2, from those allocated to placebo, 10 patients met the criteria for responders and the other 10 were classified as non-responders. From those allocated to duloxetine, eight met the criteria for responders and 11 were classified as non-responders. Patients allocated to receive placebo did not differ in baseline pain ratings when compared to those randomized to duloxetine. In both groups, responders and non-responders did not differ in baseline pain ratings, age, disease duration, depressive symptoms or medication use at the entry of the study. However, in both groups non-responders showed higher pain catastrophizing than responders. Both placebo and duloxetine produced significant reductions in pain ratings after 3-months of treatment; however, the extent of pain relief did not differ between those treated with placebo and those treated with duloxetine.

### Receptor-Enriched Analysis of Functional Connectivity by Targets (REACT)

We used the templates of the molecular density distribution of the DAT, NET, SERT and μ-opioid receptor in the REACT analysis to estimate the corresponding molecular-enriched FC maps of these systems for every subject of the two datasets. In figure 1, we provide a summary of the molecular templates (on the left) and their corresponding functional circuits (on the right) estimated by averaging the fMRI maps across healthy controls from Study 1 (for visual purposes only).

**Figure 1.**
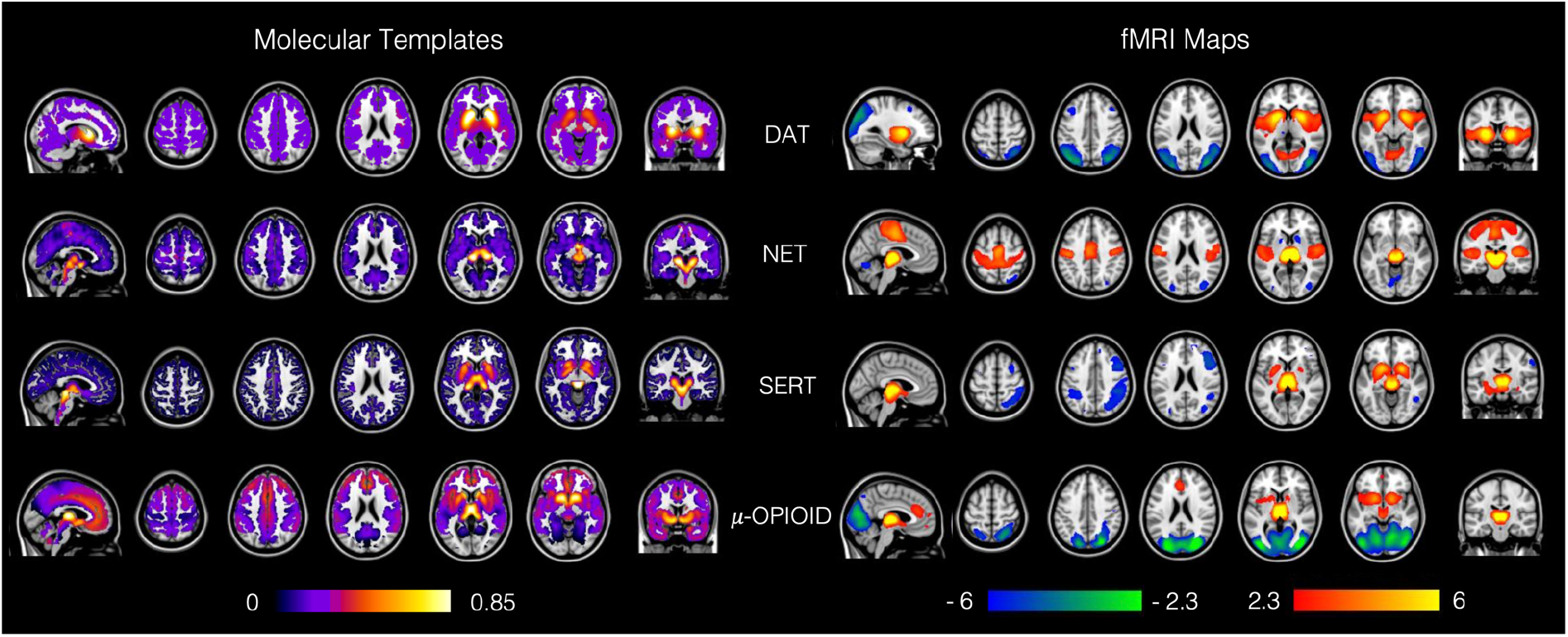
Receptor-Enriched Analysis of Functional Connectivity by Targets (REACT) multimodal framework. Maps of the molecular templates of the dopamine, noradrenaline and serotonin transporters (DAT, NET and SERT) and the μ-opioid receptor (on the left) and their respective molecular-enriched fMRI maps (on the right). The colour bar on the left represents the molecular density distribution of each template, normalised between 0 and 1 after removing either the cerebellum (for the SERT) or the occipital regions (for the NET, DAT and μ-opioid receptor) as they were used as references for quantification of the molecular data in the kinetic models for the radioligands. The colour bar on the right represents the functional connectivity of each network, expressed in z-score. The fMRI maps are averaged across the sub-set of healthy subjects from Study 1.

### NET- and SERT-related functional connectivity differs between patients with chronic knee OA and healthy controls

We investigated our first main research question by comparing the FC associated with each neurotransmitter system between OA patients and healthy controls. We started by running exploratory whole-brain two-sample t-tests (controlling for age and gender) comparing, for each neurotransmitter system, the neurotransmission-enriched FC maps of healthy controls (HC) and patients with chronic knee osteoarthritis from Study 1 (OA_1_). We found significant differences in NET and SERT-enriched FC (p_FWE_ < 0.05) between the OA_1_ and HC groups (Figure 2, panel A). Specifically, the OA_1_ group showed increased NET-enriched FC (p_FWE_ = 0.012) in a set of regions spanning the right superior and middle frontal gyrus and the frontal pole, and increased SERT-related FC (p_FWE_ = 0.032) in the frontal pole, middle and superior frontal gyrus, paracingulate gyrus and frontal medial cortex. We also found decreased SERT-enriched FC in the OA_1_ group (p_FWE_ = 0.002) in the superior and middle temporal gyrus, supramarginal gyrus and angular gyrus. Of note, only the increase in the SERT-enriched FC survived Bonferroni correction for multiple comparisons across maps and contrasts. We did not find any group differences in the DAT- and μ-opioid-enriched FC.

**Figure 2.**
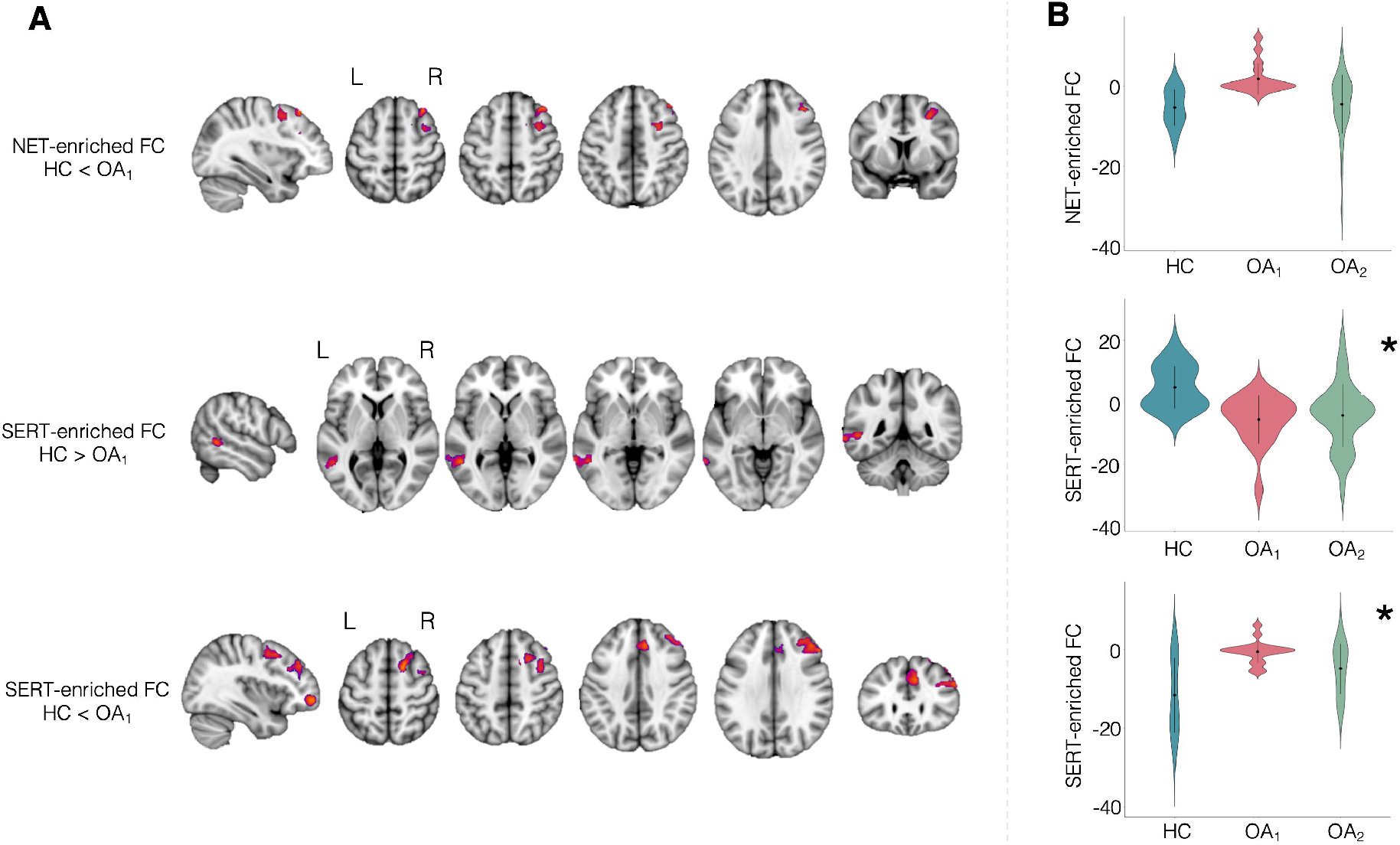
Alterations in NET- and SERT-enriched functional connectivity (FC) in patients with chronic knee osteoarthritis (OA) as compared to healthy controls (HC). (**A**) Whole-brain exploratory analysis on data from Study 1, which identified regions with significantly higher NET- and SERT-enriched FC (top and bottom rows), and other regions with reduced FC in the SERT-enriched functional maps (central row) in OA_1_ patients, as compared to healthy controls. A cluster was deemed significant if it survived p_FWE_<0.05, after correction for multiple comparisons by using the null distribution of the maximum cluster size across the image. (**B**) Hypothesis-driven analyses on extracted data from patients in Study 2 (OA_2_) showed a similar pattern of alterations in SERT-enriched FC across the two cohorts. The asterisk denotes significant differences between OA_2_ and HC (* p<0.05). Abbreviations: NET – Noradrenaline transporter; SERT – Serotonin transporter.

To validate these findings, we conducted hypothesis-driven analyses comparing SERT- and NET-enriched FC between chronic knee osteoarthritis from Study 2 (OA_2_) and HC. We extracted the mean SERT- and NET-enriched FC values in the clusters where we found case-control differences in OA_1_ from the OA_2_ patients and HC and performed direct comparisons between the two groups. The two-sample t-tests, corrected for age and gender, showed significant differences between OA_2_ and HC in SERT-related FC (HC > OA_2_: F(3,55)=7.117, p<0.0005; HC < OA_2_: F(3,55)=5.953, p=0.001), which were similar in direction and magnitude to those observed when we compared OA_1_ and HC. This analysis identified a similar pattern of alterations in SERT-enriched FC across the two cohorts of OA patients (Figure 2, panel B).

Beyond the case-control group differences, we also investigated whether NET- and SERT-enriched FC in these regions would also be able to predict the pain ratings (VAS) at entry in each of the two groups of patients. We did not find a consistent pattern of association between NET- and SERT-enriched FC and pain intensity (Supplementary Table S1).

### Placebo responders differ from non-responders in pre-treatment DAT-enriched functional connectivity

To test whether placebo responders and non-responder differ in pre-treatment FC associated with any of the neurotransmitter systems we tested, we ran exploratory whole-brain two-sample t-tests comparing FC related to each system between placebo responders and non-responders from the OA_1_ dataset, while accounting for age and gender. We found that placebo responders, as compared to non-responders, showed significant increases in DAT-enriched FC in the central and parietal opercular cortex, Heschl’s gyrus, anterior division of the superior temporal gyrus, planum polare and planum temporale (p_FWE_ = 0.027; Figure 3A). Of note, this result would not have survived a strict Bonferroni correction for multiple comparisons across maps and contrasts. No significant differences between placebo responders and non-responders were observed for NET-, SERT- and μ-opioid-enriched FC.

**Figure 3.**
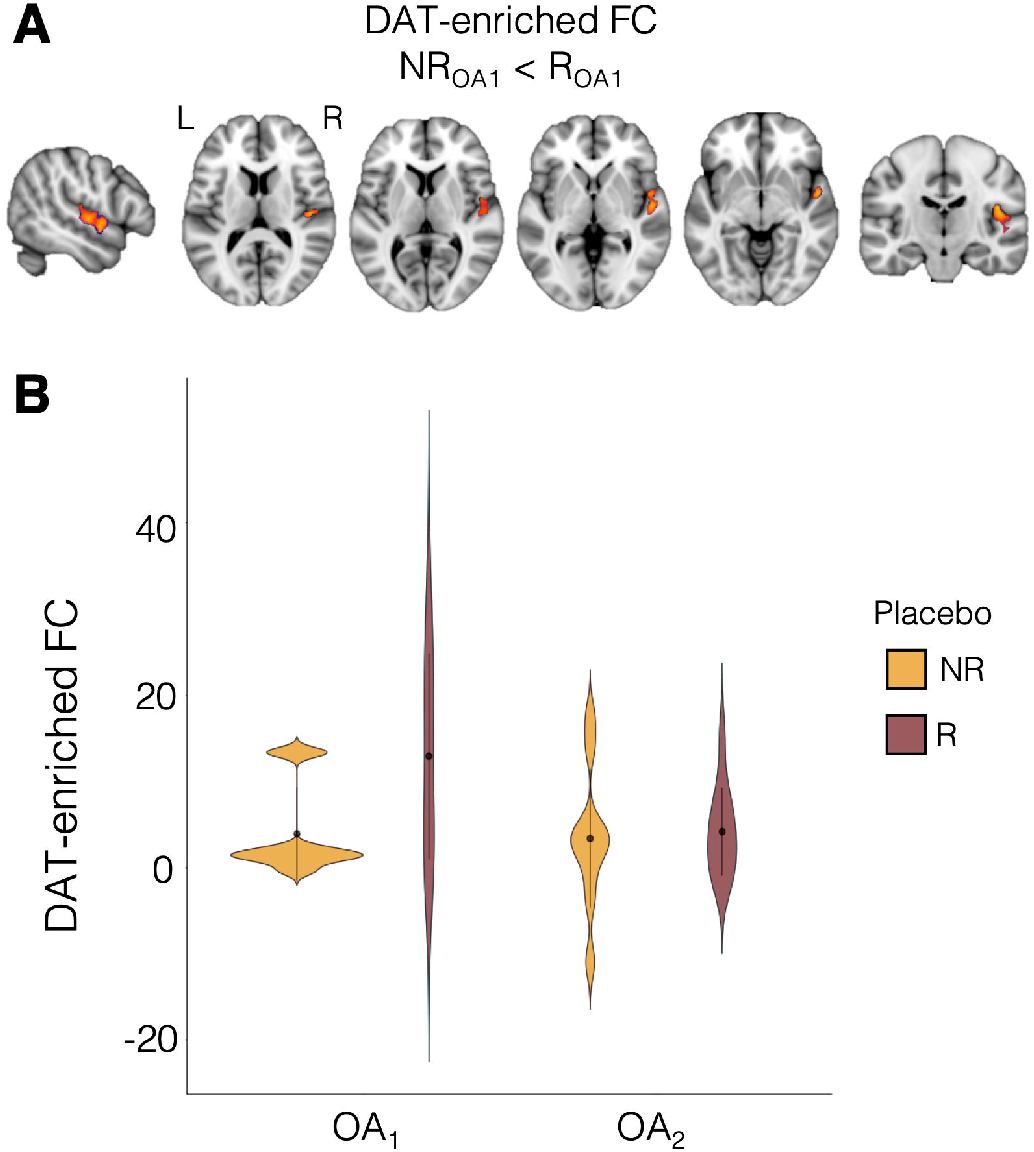
Differences in pre-treatment DAT-enriched functional connectivity (FC) between patients with chronic knee osteoarthritis (OA) who responded (R) and did not respond (NR) to placebo administration. (A) Whole-brain two-sample t-test conducted on data from Study 1 (OA_1_), which showed significantly higher pre-treatment FC in the DAT-enriched FC in placebo R as compared to NR. A cluster was deemed significant if it survived p_FWE_<0.05, after correction for multiple comparisons by using the null distribution of the maximum cluster size across the image. (B) Hypothesis-driven analysis on DAT-enriched FC values extracted from the cluster reported in (A) in patients from Study 2 did not show any significant differences between placebo R and NR. The violin plots show the mean FC values within the cluster in (A) for placebo R and NR in both Studies 1 and 2. OA_1_: N_responders_ = 8; N_non-responders_ = 9; OA_2_: N_responders_ = 10; N_non-responders_ = 10. Abbreviations: DAT – Dopamine transporter.

We performed receiver operating curve (ROC) analysis to quantify the performance of DAT-enriched FC extracted from the cluster described above to discriminate between placebo responders and non-responders in our sample. For a cut-off of 4.47 arbitrary units, DAT-enriched connectivity discriminated between responders and non-responders in our sample with a sensitivity of 75%, specificity of 77.78%, positive and negative predictive values of 75% and 77.78%, respectively (AUC = 0.722) (Supplementary Figure S1).

As a final check, we examined whether DAT-enriched connectivity differences in OA_1_ could reflect a regression to the mean phenomenon (rather than a placebo pill response) by testing whether DAT-enriched connectivity predicts symptom severity at time of entry into the study using both frequentist and Bayesian Pearson correlations. We found that DAT-enriched connectivity was not correlated with VAS prior to treatment (Supplementary Table S2).

Next, we attempted to replicate these DAT-enriched FC findings in a hypothesis-driven analysis using data from OA_2_. We extracted the mean DAT-enriched FC values in the cluster that showed a significant difference between placebo responders and non-responders in OA_1_.We then used these values in a two-sample t-test comparing placebo responders and non-responders in OA_2_, but did not find any significant group differences (Figure 3B). In a Bayesian two-sample t-test of the same data we found that, given the data, the null hypothesis was about 2.46 times more likely than the alternative hypothesis of a group difference (BF_01_ = 2.46). Further exploratory analyses at the whole-brain level did not reveal any group differences for any of the neurotransmitter systems.

### Duloxetine responders and non-responders differ in pre-treatment NET- and SERT-related functional connectivity

Finally, we tested if different patterns of pre-treatment FC related to neurotransmission underlie differences in response to different analgesic treatments. We investigated this question using data from Study 2, which allowed us to examine differences in pre-treatment FC related to neurotransmission between responders and non-responders to placebo and duloxetine. For each functional circuit, we used two-way ANCOVA to interrogate a 2-way interaction effect between treatment type (duloxetine, placebo) and treatment response (responders, non-responders) on FC, after adjusting for patients’ age and gender. We found significant two-way interaction effects for NET- and SERT-enriched FC (respectively p_FWE_ = 0.011 and p_FWE_ = 0.024; Figure 4A). In the NET-enriched FC, this interaction spanned the frontal pole, insular cortex, middle and inferior frontal gyrus, precentral gyrus, superior and middle temporal gyrus, postcentral gyrus, supramarginal gyrus and planum temporale. Similarly, in the SERT-enriched FC the interaction spanned the frontal pole, middle and inferior frontal gyrus, precentral gyrus, postcentral gyrus, superior parietal lobule, supramarginal gyrus, lateral occipital cortex and cuneal cortex. Of note, only the result related to the NET-enriched functional circuit survived Bonferroni correction for multiple comparisons across maps and contrasts. No significant interaction effects were observed in the DAT- and μ-opioid-enriched FC maps.

**Figure 4.**
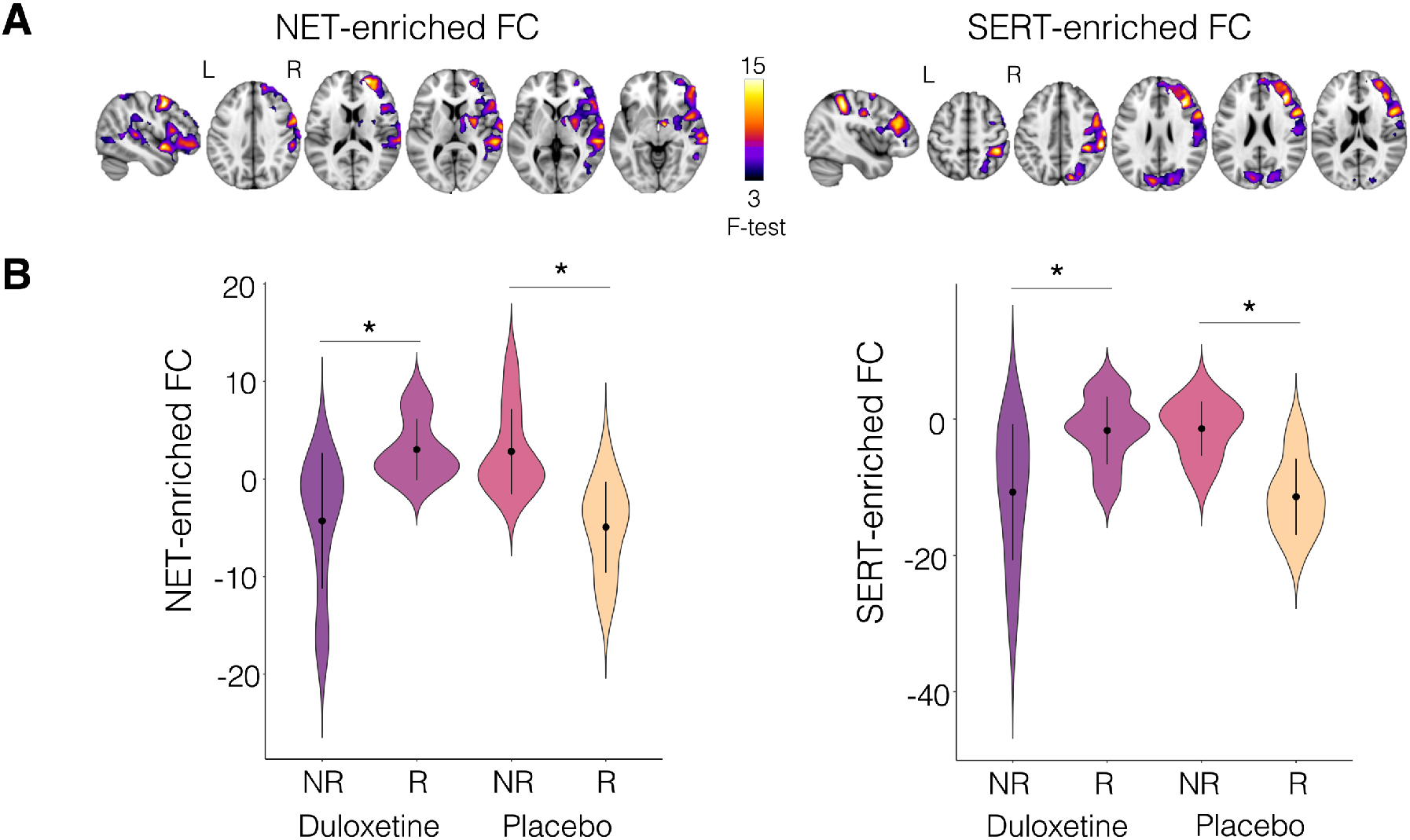
Treatment type x treatment response interaction in NET- and SERT-enriched functional connectivity (Study 2). (**A**) A two-way ANCOVA showed statistically significant interactions treatment type (duloxetine, placebo) x treatment response (responders, non-responders) in the NET- and SERT-enriched FC maps, after adjusting for patients’ age and gender. A cluster was deemed significant if it survived p_FWE_<0.05, after correction for multiple comparisons by using the null distribution of the maximum cluster size across the image. (**B**) Post-hoc tests were then run on extracted mean functional connectivity (FC) from those clusters to evaluate the simple treatment response main effects in each treatment group separately, after controlling for age and gender. Significant differences between responders (R) and non-responders (NR) are marked with an asterisk on top of the corresponding violin plots (* p<0.05 two-tailed, after Tukey correction for multiple comparison). Placebo: N_responders_ = 10, N_non-responders_ = 10; Duloxetine: N_responders_ = 8, N_non-responders_ = 11. Abbreviations: NET – Noradrenaline transporter; SERT – Serotonin transporter.

We then extracted the mean FC values within the significant clusters where we found a significant interaction in the whole-brain analyses and ran post-hoc tests to evaluate the simple main effects of response to treatment for the duloxetine and placebo groups separately (these analyses were adjusted for age and gender). In those allocated to receive duloxetine, responders had higher FC in both the NET and SERT-enriched maps than non-responders; in those allocated to receive placebo, we found the opposite (Figure 4B). The full statistics resulting from the post-hoc tests are reported in Supplementary Table S3. To investigate a potential regression to the mean phenomenon, we also used these values to calculate frequentist and Bayesian Pearson correlations between NET- and SERT-enriched FC and baseline VAS pain ratings in the placebo and duloxetine groups separately. Here, we only found a significant negative correlation SERT-enriched FC and baseline VAS in the placebo group (Supplementary Table S4).

Finally, we performed ROC analysis to quantify the performance of NET- and SERT-enriched FC extracted from the clusters described above to discriminate between responders and non-responders to duloxetine in our sample. For a cut-off of 1.45 arbitrary units, NET-enriched connectivity discriminated between duloxetine responders and non-responders in our sample with a sensitivity of 75%, specificity of 90.91%, positive and negative predictive values of 85.71% and 83.33%, respectively (AUC = 0.852). For a cut-off of −2.79 arbitrary units, SERT-enriched connectivity discriminated between duloxetine responders and non-responders in our sample with a sensitivity of 75%, specificity of 72.73%, positive and negative predictive values of 66.67% and 80%, respectively (AUC = 0.773). Direct comparisons of the AUCs of the NET- and SERT-enriched ROCs yielded no significant differences (p=0.572) (Supplementary Figure S2).

## Discussion

In this study, we investigated whether molecular-enriched FC mapping can provide insights into the brain pathophysiological mechanisms and inter-individual differences in treatment response to pharmacological analgesia during chronic pain. To that end, we applied a novel multimodal approach (REACT) to resting-state fMRI data from two studies on chronic pain patients with knee osteoarthritis. We found that, when compared to healthy controls, chronic patients with knee osteoarthritis presented alterations in the FC related to NET and SERT, two key neurotransmission systems involved in pain modulation and targeted by a class of drugs currently prescribed to control pain in these patients. Changes in SERT-enriched FC were consistent across the two datasets. In line with the known role of dopamine in expectancy and placebo response, we found that pre-treatment DAT-enriched connectivity at rest was higher in patients that responded to a 2-week period of administration of a placebo, as compared to non-responders. We also found that patients that responded to duloxetine, but not those that responded to placebo, showed higher pre-treatment NET- and SERT-enriched FC at rest, which achieved specificity and positive predictive values for future duloxetine response of up to 85.71%. Based on these findings, we suggest that molecular-enriched FC mapping might constitute a step towards mechanism-based neuroimaging biomarkers for functional alterations in chronic pain and treatment response prediction. We discuss each of our main findings below.

Our first main finding was the observation that chronic pain patients with knee osteoarthritis present alterations in NET- and SERT-enriched FC at rest, when compared to healthy controls. We validated this finding further by showing that a similar pattern of alterations was present across two datasets of patients, at least for the SERT functional circuit. This finding is interesting for two reasons. First, both the serotonin and the noradrenaline systems are part of the neurotransmission systems involved in pain control and modulation from the brain(25). The role of serotonin and noradrenaline in pain regulation is certainly complex and can encompass both inhibitory (analgesic) and excitatory (hyperalgesic) actions, depending on the site of action, cell type and type of receptor engaged(25). Yet, a vast number of studies in animal models have causally implicated alterations in serotonin and noradrenaline neurotransmission in the genesis of persistent pain (for an extensive review please see (28)). For instance, serotonin and noradrenaline depletion through repeated administration of reserpine in rodents is sufficient to induce patterns of persistent tactile allodynia and it has recently been used as a fibromyalgia-like animal model for research on disease mechanisms and drug development(29–32). Second, both the serotonin and noradrenaline transporters are targeted by drugs currently prescribed to control pain in patients with chronic pain, such as tricyclic antidepressants or non-selective inhibitors of the reuptake of serotonin and noradrenaline(20). While these drugs are often prescribed to patients with chronic pain, the understanding of the precise mechanisms through which they might reduce pain in these patients is still relatively poor(20). The animal literature is strongly supportive of the hypothesis that antidepressants might enhance the engagement of descending inhibitory pain pathways by increasing serotonin and noradrenaline neurotransmission(20) (though this picture is likely to be more complex given that, for instance, different serotonin receptors can be inhibitory or facilitatory(33), or that increases in noradrenaline in regions of the brain involved in descending pain modulation, such as the dorsal reticular nucleus, can also facilitate pain(34)). However, whether the same mechanisms are responsible for the clinical effects observed in chronic pain patients has never been explored in depth. Based on our findings we could speculate that one of the mechanisms through which these drugs might improve pain control in chronic pain patients is through normalizing the alterations in NET- and SERT-enriched FC we report here. While we could not test this hypothesis using our datasets since imaging data at follow-up were not collected for these two studies, we believe this is an interesting question for future work, as it could help to strengthen the rationale for using SERT- and NET-targeting compounds to treat chronic pain.

We did not find any alteration in DAT- or μ-opioid-enriched connectivity at rest in any of the two OA datasets we analysed. This was surprising for several reasons. First, the opioid system has a well-established role in pain regulation(35, 36) and the dopamine system has equally been suggested to be involved in the supraspinal modulation of pain(37, 38). Second, alterations in opioid and dopaminergic neurotransmission in chronic pain have been reported in human PET studies(39, 40). For instance, chronic neuropathic pain was shown to be associated with higher striatal dopamine D_2_/D_3_ receptor availability, for which low endogenous dopamine tone is a plausible explanation(41). Alterations in μ-opioid receptor availability has been showed across chronic pain conditions (39), including arthritis(42). Third, opioids figure among the pharmacological agents used to manage chronic pain(43) and can achieve effective analgesia in at least some types of chronic pain(44). Based on these lines of evidence, it would be plausible that chronic pain in OA might be associated with DAT- or μ-opioid-enriched FC changes. While we can only speculate around null findings, we believe at least two factors might have contributed to the lack of DAT- or μ-opioid-enriched connectivity we report here. First, we based our analyses on target-enriched FC measured at rest. Hence, we cannot exclude that such alterations might emerge under nociceptive stimulation, which has been shown to recruit the opioid system in human PET studies(45), and might enhance case-control differences in μ-opioid-enriched connectivity, if they exist. Second, while one of the exclusion criteria in both datasets was current treatment with MAO inhibitors or any centrally acting drug for analgesia and depression, to be eligible patients needed daily pain medication to manage symptoms. While we are unaware of the exact drug class used by these patients before enrolment, it is possible that such treatment might have mitigated potential case-control differences in μ-opioid- or DAT-enriched connectivity, if they existed. Future studies exploring these questions will be important.

Our second main finding was the observation that patients experiencing analgesia in response to a course of 2 weeks of placebo administration (Study 1) present higher pre-treatment DAT-enriched FC, but not SERT-, NET- or μ-opioid-enriched FC, than patients that did not respond. DAT-enriched FC was not related to disease burden prior to start of placebo treatment in Study 1, diminishing the possibility that the measure is related to regression to the mean rather than a true placebo response. Furthermore, this difference is also unlikely to be confounded by differences between responders and non-responders in disease duration, depressive symptoms, pain catastrophizing or medication use since the groups did not significantly differ in any of these variables (see original study(24)). This finding suggests that inter-individual differences in DAT-enriched FC might contribute to explaining why patients differ in their responses to placebo. Positive medical responses to placebo treatments are a well-recognized phenomenon observed in many pathologies, particularly for neurological and painful conditions(46, 47). Analgesia in response to placebo is widely observed in pain clinical trials, in which it often exhibits sustained effectiveness rivalling in magnitude the one from the active treatment(48, 49). Historically, the placebo effect has been thought of as the end-product of biases in subjective symptom reporting(50). This interpretation has evolved through increasing evidence that the placebo effect is mediated by specific neural mechanisms(49–51). One of the key theories around the neurobiological mechanisms underlying the placebo effect postulates that it represents a form of reward expectation processing(52). Dopamine is thought to be centrally involved in reward expectation and variations from expected outcomes (prediction errors), and has therefore been linked to placebo effects(53). For instance, one human PET study has shown that placebo-induced analgesia is associated with decreases in binding [^11^C]raclopride to D2/D3 dopamine receptors in the basal ganglia, possibly reflecting increases in the release of dopamine in these regions(27). The same study also reported placebo-induced decreases in [^11^C]carfentanil binding to the μ-opioid receptor, pointing to engagement of the endogenous opioid system during placebo-induced analgesia. However, changes in [^11^C]raclopride binding in the nucleus *accumbens* emerged as the strongest predictor of placebo-induced analgesia, accounting for 25% of the variance alone. Another study has shown that individual differences in reward response can explain placebo-induced effects and expectations(54). The differences in DAT-enriched FC between patients that responded versus those that did not respond to administration of placebo we report here are broadly compatible with this idea. Assuming that higher DAT-enriched FC might be driven by strongest dopamine-related neurotransmission within the dopaminergic circuits (which we index here through DAT density distribution in the brain), then it is plausible that those patients with strongest dopamine-related neurotransmission might benefit the most from expectancy effects, which rely on dopamine release and are at the core of the placebo effect.

This finding was nevertheless not replicated in the hypothesis-driven analysis on data from Study 2, where placebo and non-placebo responders did not differ in pre-treatment DAT-enriched FC. The lack of between-groups differences was supported by our Bayesian analysis where we found the null hypothesis to be 2.46 times more likely given the data than the alternative hypothesis. We should highlight though that there is at least one important methodological difference between Study 1 and 2 that could potentially account for this discrepancy. In Study 1, placebo response was evaluated after 2 weeks of placebo administration, while in study 2 the placebo protocol lasted for 3 months. Studies have shown that duration of administration is a determinant of placebo response(55, 56). Hence, it is possible that while inter-individual differences in DAT-enriched FC are particularly relevant to explain differences in short-term response to placebo, other mechanisms might be involved in the long term. Until further larger studies will revisit these findings, we urge for some caution when interpreting this association between pre-treatment DAT-enriched FC and placebo response. We also note the lack of differences between placebo responders and non-responder on μ-opioid-enriched FC in this study, despite previous evidence that the endogenous opioid system is recruited during placebo-induced analgesia(27). However, as explained above, whether that might reflect the fact that FC was measured at rest or that some carry-over effects of previous analgesic treatments biased this result is unclear.

Our third key finding was the observation that patients that responded to a course of treatment with duloxetine showed higher pre-treatment NET- and SERT-enriched FC (but not DAT or μ-opioid-enriched FC) than those that did not respond, while in those patients allocated to placebo we observed the opposite trend – i.e., lower pre-treatment NET- and SERT-related FC. Furthermore, NET- and SERT-related FC discriminated between duloxetine treatment response groups in our sample with good sensitivity and positive predictive values of up to 85.71%. NET-enriched FC was not related to disease burden prior to start of placebo or duloxetine treatment in Study 2, diminishing the possibility that the measure is related to regression to the mean rather than a true response. SERT-enriched FC was also not related to baseline pain ratings in the duloxetine group, but we did find a significant negative correlation in those patients allocated to receive placebo. Furthermore, these differences are unlikely to be driven by differences between responders and non-responders within each group in disease duration, depressive symptoms or medication use, since the groups did not significantly differ in any of these variables (see original study(24)). Altogether, these findings suggest that pre-treatment NET- and SERT-related FC measured at rest might hold promise as a biomarker for duloxetine analgesia response in OA chronic pain patients.

Duloxetine is a non-selective inhibitor of the reuptake of serotonin and noradrenaline, which enhances their bioavailability at the synaptic level(20). The main mechanisms suggested to underlie the analgesic effect observed under duloxetine include enhancement of descending inhibitory pain pathways from the brain through potentiation of serotoninergic and noradrenergic transmission, with consequent inhibition of ascendant transmission of nociceptive inputs from the spinal cord(20) (although peripheral actions have also received support from some preclinical studies(57)). This mechanism has received indirect support from a previous study linking response to duloxetine in painful diabetic neuropathy to the integrity of the descending pain inhibitory pathways, as assessed by conditioned pain modulation(58). Therefore, the fact that only pre-treatment differences in SERT and NET-enriched FC exist between duloxetine responders and non-responders matches the pharmacodynamics of the drug and aligns with the basic drug mechanisms through which most likely it induces analgesia. Based on this observation, we suggest that target-enriched FC mapping might open a new avenue in neuroimaging biomarkers of pharmacological treatment response in chronic pain, bringing the advantage of allowing to establish a clearer mechanistic link between the neuroimaging biomarker being measured and the neurotransmission-related mechanisms through which a pharmacological treatment targeting a specific neurochemical system might induce analgesia. For instance, since duloxetine acts by inhibiting the reuptake of serotonin and noradrenaline, it is conceivable that its ability to enhance serotoninergic or noradrenergic transmission is moderated by the amounts of these neurotransmitters available in the synapses, promoting lower accumulation of serotonin/noradrenaline in those where synthesis capacity and tonic release is reduced. Following this line of thought, patients with lower bioavailability of serotonin/noradrenaline, which for this reason might show lower SERT- and NET-related FC connectivity, might not benefit from duloxetine treatment as much. We should highlight though that at this point any relationship between bioavailability of specific neurotransmitters and target-enriched FC remains speculative and will require validation in further studies.

While we could not find significant differences between NET- and SERT-enriched FC ability to discriminate between responders and non-responder to duloxetine, we note that NET-enriched FC showed higher AUC, specificity and positive predictive value. This empirical observation matches the clinical evidence that non-selective inhibitors of the reuptake of serotonin and noradrenaline might be superior to selective inhibitors of the reuptake of serotonin in promoting analgesia in patients with chronic pain(20, 59). This superiority is thought to be linked to the fact that antidepressants which also increase the levels of noradrenaline by inhibiting NET might block the spinal transmission of nociceptive input directly through acting on spinal α2 receptors(20).

Our study has some limitations worth mentioning. First, although REACT improves the specificity of FC analysis, the approach remains relatively indirect and relies on molecular templates estimated in independent cohorts of healthy individuals. Therefore, further specification from intra-regional variation across patients is not possible using the current dataset as it would require PET data for each ligand and patient. The availability of PET data from the same cohort of patients would allow the creation of patient-specific templates, which might enhance the accuracy of the maps of FC related to each target. This should be examined in future studies validating our work further. Second, the sample size of both cohorts was relatively small. This was in part mitigated by the fact that we attempted to replicate some of our findings in Study 1 using a second cohort of patients from study 2. Nevertheless, future larger studies revisiting these findings will be welcome. Third, we explored the ability of NET- and SER-enriched FC to discriminate between responders and non-responders to duloxetine and provided preliminary evidence that it might offer good specificity and true positive predictive value in discriminating between them in our sample. However, given that we conducted such analyses using data from the same sample we used for biomarker discovery, these results cannot reiterate the validity of our candidate biomarkers in predicting placebo or duloxetine response yet. The validity of our candidate biomarkers will need to be inspected in future studies using larger independent cohorts. Finally, our findings are restricted to OA patients and to the prediction of placebo and duloxetine response in this group of chronic pain patients; hence, direct extrapolation to other chronic pain conditions or other pharmacological analgesics should be avoided. Indeed, chronic pain manifests in a range of clinical phenotypes; and even within the boundaries of a specific chronic pain syndrome such as OA, it is likely that different pathophysiological mechanisms are in play in different patients(60, 61). However, from the findings we gathered in this study, we suggest that future studies expanding the approach we presented here to other chronic pain populations and drug classes are worth investing and might reveal fruitful.

In conclusion, while further clinical validation in larger cohorts is warranted, we suggest that molecular-enriched FC mapping might hold promise as a new mechanistic-informed biomarker for functional brain alterations and prediction of response to pharmacological analgesia in chronic pain. The mechanistic insights introduced by this approach might help to identify chronic pain mechanisms enabling rational and individualized treatment choice. Ultimately, these informed decisions might contribute to decrease unnecessary exposures of patients to ineffective therapies and undesirable side-effects, facilitate treatment adherence and accelerate pain control without long periods of treatment trial-and-error, decreasing the chance that the pain becomes intractable(62).

## Methods

### Participants and study design

For this work, we used two openly available rs-fMRI datasets^(24)^ of chronic knee OA pain patients who underwent pre-treatment brain scans in two clinical trials. The full details on demographics, inclusion and exclusion criteria have been provided in the original article(24). Here, we will simply present a brief summary to help to contextualize the reader. Study 1 was a 2-week single-blinded placebo pill trial where 17 OA patients (M/F:8/9; 56.9 ± 5.7 years) ingested a lactose placebo pill once a day for 2 weeks. Study 2 was a 3-month double-blinded randomized trial in which 39 OA patients ingested either placebo pills (n = 20; M/F: 9/12; 57.6 ± 9.5 years) or duloxetine (n = 19; M/F: 9/10; 59.2 ± 4.6 years) at a dose of 30 mg for the first week and escalated to 60 mg for the rest of the treatment period, except for the last week, when the dose was decreased back to 30 mg. In addition, Study 1 also included 20 age-matched healthy control subjects (M/F:10/10; 57.9 ± 6.7 years). For Studies 1 and 2, behavioural and clinical parameters were obtained before and after treatment, while brain scans were collected only before treatment. Patients were asked to discontinue their medications 2 weeks before the beginning of the trial and were provided with acetaminophen as rescue medication. All participants gave written informed consent to procedures approved by the Northwestern University Institutional Review Board committee (STU00039556).

### Behavioural and clinical measures

Patients from both studies completed a general health questionnaire, a Visual Analogue Scale (VAS) on a 0 to 10 scale for their knee OA pain, the Western Ontario and McMaster Universities Osteoarthritis Index (WOMAC), the Beck Depression Inventory (BDI) and the Pain Catastrophizing Scale (PCS) (please note we could only access the raw VAS and WOMAC data). All questionnaires were administered on the day of brain scanning. In Study 2, to partially compensate for regression to the mean effects, VAS was measured 3 times over a 2-week period prior to the start of treatment and after cessation of medication use, averaged, and used as baseline. Analgesic response was defined a priori on an individual basis as at least a 20% decrease in VAS pain from baseline to the end of treatment period; otherwise, subjects were classified as non-responders. This threshold for analgesic response was chosen following the same procedure adopted by the previous study on these datasets(24). While the choice of this threshold is arbitrary, we note though that a 20% reduction in VAS ratings of pain is higher than the 15% considered to be minimal clinically important(63). As a further reference, a 30% reduction in VAS ratings of pain is typically considered a clinical important pain diminution(64).

### Image acquisition

For all participants in Studies 1 and 2, imaging data were collected with a 3T Siemens Trio whole-body scanner. A 3D T1-weighted anatomical scan was obtained for each participant using an MPRAGE acquisition (voxel size = 1×1×1 mm; TR/TE = 2,500/3.36 ms; flip angle = 9°; in-plane matrix resolution = 256 × 256; slices = 160; field of view = 256 mm). Functional MRI data were obtained during rest using a multi-slice T2*-weighted echo-planar sequence (TR/TE = 2500/30 ms, flip angle = 90°, number of slices = 40, slice thickness = 3 mm, and in-plane resolution = 64 × 64; number of volumes = 300). All MRI data are available on https://openneuro.org/datasets/ds000208/versions/1.0.0.

### Image pre-processing

The rs-fMRI datasets from Studies 1 and 2 were pre-processed using FMRIB Software Library (FSL). Pre-processing steps included volume re-alignment with MCFLIRT(65), non-brain tissue removal with the brain extraction tool (BET)(66), an initial spatial smoothing with a 6-mm FWHM Gaussian kernel and de-noising with ICA-based Automatic Removal Of Motion Artifacts (ICA-AROMA)(67). Additionally, subject-specific WM and CSF masks, obtained from the segmentation of the subjects’ structural images and eroded in order to minimize the contribution of grey matter partial volume effects, were used to extract and regress out the mean WM and CSF signals from each participant’s pre-processed dataset. A high-pass temporal filter with a cut-off frequency of 0.005 Hz was applied, followed by an additional spatial smoothing at FWHM = 6mm, in order to obtain a final smoothing of the fMRI images of approximately 8mm (FWHM^2^_final_ = FWHM^2^_1_+FWHM^2^_2_). A study-specific template representing the average T1-weighted anatomical image across subjects was built using the Advanced Normalization Tools (ANTs)(68). Each participant’s dataset was co-registered to its corresponding structural scan, then normalized to the study-specific template before warping to standard MNI152 space. Images were finally resampled at 2 mm^3^ resolution.

### Functional connectivity analysis with REACT

For the analysis with REACT, we used the templates of the molecular density distribution of the DAT, NET, SERT and μ-opioid receptor. The DAT map is a publicly available template of 123I-Ioflupane SPECT images (https://www.nitrc.org/projects/spmtemplates) from 30 HC without evidence of nigrostriatal degeneration(69). The NET atlas is a publicly available template of the [^11^C]MRB PET brain parametric maps from 10 HC (M/F: 6/4; 33.3 ± 10 years)(70, 71). The SERT atlas is a publicly available template(72) of [^11^C]DASB PET images of 210 healthy controls from the Cimbi database(73). The μ-opioid receptor map is a publicly available template of [^11^C]Carfentanil PET images of 89 HC (https://identifiers.org/neurovault.image:115126). All molecular atlases were normalised by scaling the image values between 0 and 1, although preserving the original intensity distribution of the images, and masked using a standard grey matter mask. Of note, for each atlas we masked out the regions that were used as references for quantification of the molecular data in the kinetic models for the radioligands, namely the occipital areas for DAT, NET and μ-opioid receptor, and the cerebellum for SERT. Finally, we resampled the SERT image in order to have all atlases in standard MNI space with 2 mm^3^ voxel size.

A detailed explanation of REACT methodology and its applications can be found elsewhere(22, 23). In brief, the functional circuits related to the DAT, NET SERT and μ-opioid receptor systems were estimated using a two-step multivariate regression analysis(74, 75) implemented with the *fsl_glm* command of FSL. In the first step, the rs-fMRI volumes were masked using a binarized atlas derived from the molecular data to restrict the analysis to the voxels for which the density information of the neurotransmitter was available in the template. Then, the molecular templates were used as a set of spatial regressors to weight the rs-fMRI images and estimate the dominant BOLD fluctuation related to each molecular system at the subject level. Those subject-specific time series were then used as temporal regressors in a second multivariate regression analysis to estimate the subject-specific spatial map associated with each molecular atlas. The output consists of four maps per subject, each one reflecting the molecular-enriched FC associated with a specific neurotransmitter. At this stage, the analysis was conducted on the whole grey matter volume. Both data and the design matrix were demeaned (--demean option); the design matrix columns were also normalised to unit standard deviation with the --des_norm option(74).

### Statistics

For our analyses on case-control differences, first we run exploratory whole-brain two-sample t-tests comparing OA_1_ and healthy controls, after controlling for age and gender, for each neurotransmission system separately. For this and all subsequent whole-brain analyses, we applied cluster-based inference within Randomise(76), using 5,000 permutations per test and contrast, considering a cluster significant if p_FWE_ < 0.05, corrected for multiple comparisons by using the null distribution of the maximum cluster size across the image. Then, we used a two-sample t-test in SPSS (version 27) to investigate differences in mean SERT- and NET-enriched FC values, extracted from the clusters where we found significant OA_1_ vs healthy controls differences, between OA_2_ and healthy controls after controlling for age and gender. For our exploratory analyses on differences in pre-treatment FC between placebo responders and non-responders, we used whole-brain two-sample t-tests comparing responders and non-responders, after controlling for age and gender, for each neurotransmission system separately. Our hypothesis-driven analysis for DAT-enriched differences in pre-treatment FC between responders and non-responders in Study 2 used a two-sample t-test implemented in SPSS. We investigated two-way interactions between treatment type (placebo, duloxetine) and treatment response (responders, non-responders) using exploratory whole-brain F-tests, adjusting for age and gender, for each neurotransmission system separately. We followed up each significant interaction by extracting the mean FC values within the significant clusters found in the whole-brain interaction analysis and ran post-hoc tests in SPSS to evaluate the simple main effects of treatment response within each treatment type groups separately, after controlling for age and gender. Levene’s test was also performed to check the homogeneity of variances. The correlations between mean molecular-enriched FC and pain ratings were calculated using Pearson’s correlations (bootstrapping 1,000 samples), as implemented in SPSS. The ROC analyses on treatment response discrimination were implemented in JAMOVI, using the PPDA package. Here, we defined response as a binary variable taking the value 0 for non-responders and 1 for responders. We set 1 (responders) as the positive class and used the Youden’s J statistic to select the cut-off that maximizes the performance of the discrimination (larger is better). All Bayesian analyses were implemented in JAMOVI (version 1.16.2.0), using the default uninformative priors from the software. An increase in Bayes Factor (BF) in our analyses corresponds to an increase in evidence in favour of the null hypothesis. To interpret BF, we used the Lee and Wagenmakers’ classification scheme(77): BF < 1/10, strong evidence for alternative hypothesis; 1/10<BF<1/3, moderate evidence for alternative hypothesis; 1/3 < BF < 1, anecdotal evidence for alternative hypothesis; BF > 1, anecdotal evidence for the null hypothesis; 3<BF<10, moderate evidence for the null hypothesis; BF > 10, strong evidence for the null hypothesis.

## Author contributions

DM and OD designed the study, analysed the data and drafted the first version of the manuscript. All authors contributed to the interpretation of the findings and provided critical revisions to the final version of the manuscript. All authors approved the final version of the manuscript.

## Acknowledgements

We would like to thank the authors of the original study for making the data available and all patients contributing data to this study. We would also like to thank Prof. Swen Hesse, Prof. Osama Sabri and Dr Michael Rullmann (Department of Nuclear Medicine, University of Leipzig, Leipzig, Germany; Integrated Research and Treatment Center Adiposity Diseases, Leipzig University Medical Center, Leipzig, Germany) for kindly providing the PET template of the noradrenaline transporter.

We would like to thank also the National Institute for Health Research (NIHR) Biomedical Research Centre at South London and Maudsley NHS Foundation Trust and King’s College London for their ongoing support of our research. The views expressed are those of the authors and not necessarily those of the NHS, the NIHR or the Department of Health and Social Care.

