## Supplementary material for "A candidate neuroimaging biomarker for detection of neurotransmission-related functional alterations and prediction of pharmacological analgesic response in chronic pain"

**Supplementary Table S1. Correlations between NET- and SERT-enriched functional connectivity and pain burden.** We calculated frequentist and Bayesian Pearson correlations between mean NET and SERT-enriched FC values extracted from the clusters where we found differences between patients and healthy controls in Study 1 and baseline pain ratings. We repeated the same procedure for data extracted from patients in Study 2. Statistical significance was set at $p < 0.05$ (two-tailed). Abbreviations: NET – Noradrenaline transporter; SERT – Serotonin transporter; VAS – Visual analogue scale.

| | **NET-enriched FC ($HC < OA_1$)** | | | **SERT-enriched FC ($HC > OA_1$)** | | | **SERT-enriched FC ($HC < OA_1$)** | | |
|---|---|---|---|---|---|---|---|---|---|
| Baseline VAS | **r** | **p-value** | **BF01** | **r** | **p-value** | **BF01** | **r** | **p-value** | **BF01** |
| $OA_1$ | 0.227 | 0.381 | 2.34 | 0.170 | 0.514 | 2.74 | -0.001 | 0.998 | 3.34 |
| $OA_2$ | 0.225 | 0.096 | 0.881 | 0.323 | 0.015 | 0.468 | 0.227 | 0.092 | 0.708 |

**Supplementary Figure S1. Performance of DAT-enriched functional connectivity in discriminating between placebo responders and non-responders.** Here, we present simply the Receiver Operating Curve; full performance metrics reported in the main manuscript. Abbreviations: DAT – Dopamine transporter; FC – Functional Connectivity.

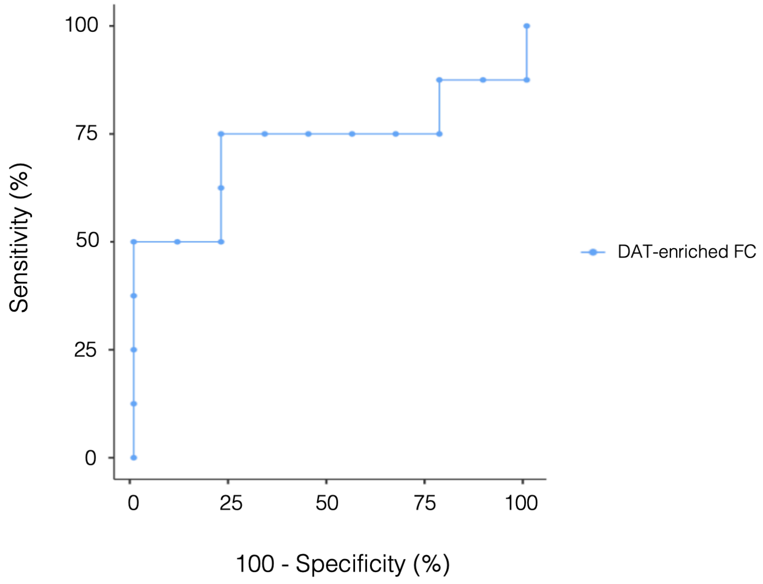


**Supplementary Table S2. Correlations between pre-treatment DAT-enriched functional connectivity (FC) and baseline visual analogue scale (VAS) rating of pain in patients from Study 1.** With this analysis, we aimed to examine whether the differences we detected in DAT-enriched FC between placebo responders and non-responders could simply reflect a regression to the mean phenomenon. We calculated frequentist and Bayesian pearson correlations between mean DAT-enriched FC values extracted from the cluster where we found differences between placebo responders and non-responders in the exploratory whole-brain analysis and baseline VAS pain ratings. Statistical significance was set at $p < 0.05$ (two-tailed). Abbreviations: DAT – Dopamine transporter; FC – Functional Connectivity

| | DAT-enriched FC | | |
|---|---|---|---|
| **Study 1** | **r** | **p-value** | **BF01** |
| Baseline VAS | -0.025 | 0.923 | 3.32 |

**Supplementary Table S3. Simple main effects of treatment response in the duloxetine and placebo groups**. We used a general linear model, considering treatment response (responders, non-responders) as a fixed factor and age and gender as covariates. Statistical significance was set at $p < 0.05$ (two-tailed), after correction for multiple comparisons with Tukey. Abbreviations: NET – Noradrenaline transporter; SERT – Serotonin transporter; FC – Functional Connectivity.

| | **NET-enriched FC** | | | **SERT-enriched FC** | | |
|---|---|---|---|---|---|---|
| **Treatment group** | **F** | **p-value** | **Partial $\eta^2$** | **F** | **p-value** | **Partial $\eta^2$** |
| Duloxetine | $F(1,15) = 3.76$ | 0.034 | 0.43 | $F(1,15) = 4.60$ | 0.018 | 0.479 |
| Placebo | $F(1,16) = 4.45$ | 0.019 | 0.455 | $F(1,16) = 8.57$ | 0.001 | 0.616 |

**Supplementary Table S4. Correlations between pre-treatment SERT and NET-enriched functional connectivity (FC) and baseline visual analogue scale (VAS) rating of pain in patients from Study 2.** With this analysis, we aimed to examine whether the differential differences we detected in NET and SERT-enriched FC between placebo and duloxetine responders and non-responders could simply reflect a regression to the mean phenomenon. We calculated frequentist and Bayesian Pearson correlations between mean NET and SERT-enriched FC values extracted from the clusters where we found significant interactions between treatment type and treatment response in the exploratory whole-brain analyses and baseline VAS pain ratings, for each treatment type group separately. Statistical significance was set at $p < 0.05$ (two-tailed). Abbreviations: NET – Noradrenaline transporter; SERT – Serotonin transporter; FC – Functional Connectivity; VAS – Visual analogue scale; BF – Bayes Factor.

| | NET-enriched FC | | | SERT-enriched FC | | |
|---|---|---|---|---|---|---|
| **Study 2** | **r** | **p-value** | **BF01** | **r** | **p-value** | **BF01** |
| Baseline VAS (Placebo group) | -0.425 | 0.062 | 0.707 | -0.582 | 0.007 | 0.123 |
| Baseline VAS (Duloxetine group) | -0.287 | 0.233 | 1.81 | 0.027 | 0.914 | 3.50 |

**Supplementary Figure S2. Performance of NET- and SERT-enriched functional connectivity in discriminating between duloxetine responders and non-responders.** Here, we present simply the Receiver Operating Curve; full performance metrics reported in the main manuscript. Abbreviations: NET – Noradrenaline transporter; SERT – Serotonin transporter; FC – Functional Connectivity.

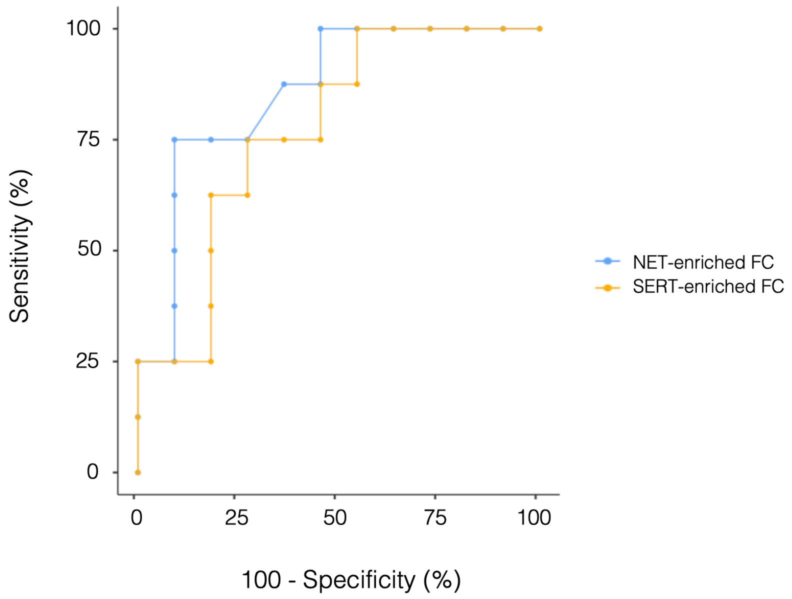
